# Anatomical circuits for flexible spatial mapping by single neurons in posterior parietal cortex

**DOI:** 10.1101/2024.04.23.590686

**Authors:** Bashir Ahmed, Hee Kyoung Ko, Maria Rüsseler, Jackson E. T. Smith, Kristine Krug

## Abstract

Primate lateral intraparietal area (LIP) is critical for cognitive processing. Its contribution to categorization and decision-making has been causally linked to neurons’ spatial sensorimotor selectivity. We reveal the intrinsic anatomical circuits and neuronal responses within LIP that provide the substrate for this flexible generation of motor responses to sensory targets. Retrograde tracers delineate a loop between two distinct operational compartments, with a sensory-like, point-to-point projection from ventral to dorsal LIP and an asymmetric, more widespread projection in reverse. Neurophysiological recordings demonstrate that especially more ventral LIP neurons exhibit motor response fields that are spatially distinct from its sensory receptive field. The different associations of response and receptive fields in single neurons tile visual space. These anatomical circuits and neuronal responses provide the basis for the flexible allocation of attention and motor responses to salient or instructive visual input across the visual field.

## Introduction

Posterior Parietal Cortex (PPC) constitutes a hub of brain areas which compute sensorimotor transformations and contribute to cognitive functions like action selection, spatial awareness, sensory evaluation and directing attention in primates [1-3]. One of the most studied PPC areas is lateral intraparietal area (LIP), which has been afforded central roles in attentional processing, decision-making, visual categorisation and oculo-motor planning [4-8]. Cyto-architectonical and myelin-architectural studies have shown that LIP can be subdivided into a dorsal (LIPd) and ventral part (LIPv) [9, 10]. The connectivity with other cortical areas differs between LIPd and LIPv [9, 11-14]. LIPv provides strong feed-forward connections to oculomotor areas such as the frontal eye fields (FEF) and the superior colliculus (SC) as well as receiving input (amongst others) from dorsal visual area V5/MT, a connection missing for LIPd. In contrast, LIPd receives input from ventral stream visual areas and has unique connections with the cingulate cortex and the dysgranular insular cortex. The differential profile of laminar terminations of projections from area FEF shows that LIPd receives feedforward input from FEF while LIPv receives feedback projections [15]. These findings taken together with inactivation studies suggest that LIPd and LIPv should also differ in their functional roles [16]. Despite these differences, the intrinsic circuitry within LIP remains unclear.

Area LIP lies on the posterior-lateral bank of the intraparietal sulcus. From the earliest neurophysiological studies, neurons in area LIP have been characterized by their “motor field” (MF), often also termed “response field”, which signals the direction and amplitude of an upcoming saccades into the MF [17], even in the absence of a visual target stimulus [18, 19]. In other studies, LIP neurons were shown to respond to visual stimuli in their sensory “receptive field” (RF) [20] and can be tuned to features like direction of motion or binocular disparity [21, 22]; RFs are re-mapped to their expected location *before* an upcoming saccade and RFs are also modulated by their behavioural significance [3, 23]. Visual RFs have usually been described as co-localised with MFs and to represent information about saccade targets [19, 24]. Our results presented here show that this is one option among a range of actual spatial configurations.

Due to its strong activation by visual stimuli and planned saccades, it is not surprising that LIP is crudely topographically organized, at least in LIPv, with a preference for the contralateral visual field [9, 25, 26]. The type of neuronal responses and their reported organisation tend to differ depending on whether the subject carries out a (saccade) task or was anaesthetized and related to this, how initial screening for neurons was conducted (during saccades or by passively viewing visual stimuli), such that two intersecting topographic maps have been suggested [27]. The alignment between the two maps and especially whether they intersect in single neurons, which is critical to how LIP neurons compute sensorimotor transformations, is one of the questions we address in this study.

## Results

### LIPv neurons send point-to-point projection to a single site in LIPd

To investigate the intrinsic connectivity of cortical area LIP, we placed single small, focal injections of retrograde tracers Choleratoxin b (CTb) or Fluorogold (FG) into LIP in one hemisphere of six Rhesus macaques (*macaca mulatta*) (Fig. 1). After 2-12 days survival, animals were transcardially perfused and tissue histologically processed. A one-in-five series of parasagittal sections (50 μm) of the injected hemisphere were stained for myelin to ascertain LIPv and LIPd borders (Fig. 1B). Retrogradely labelled cell bodies within LIP were analysed using Neurolucida in another one-in-five series of the same hemisphere (Fig. 1C,E).

**Fig. 1.**
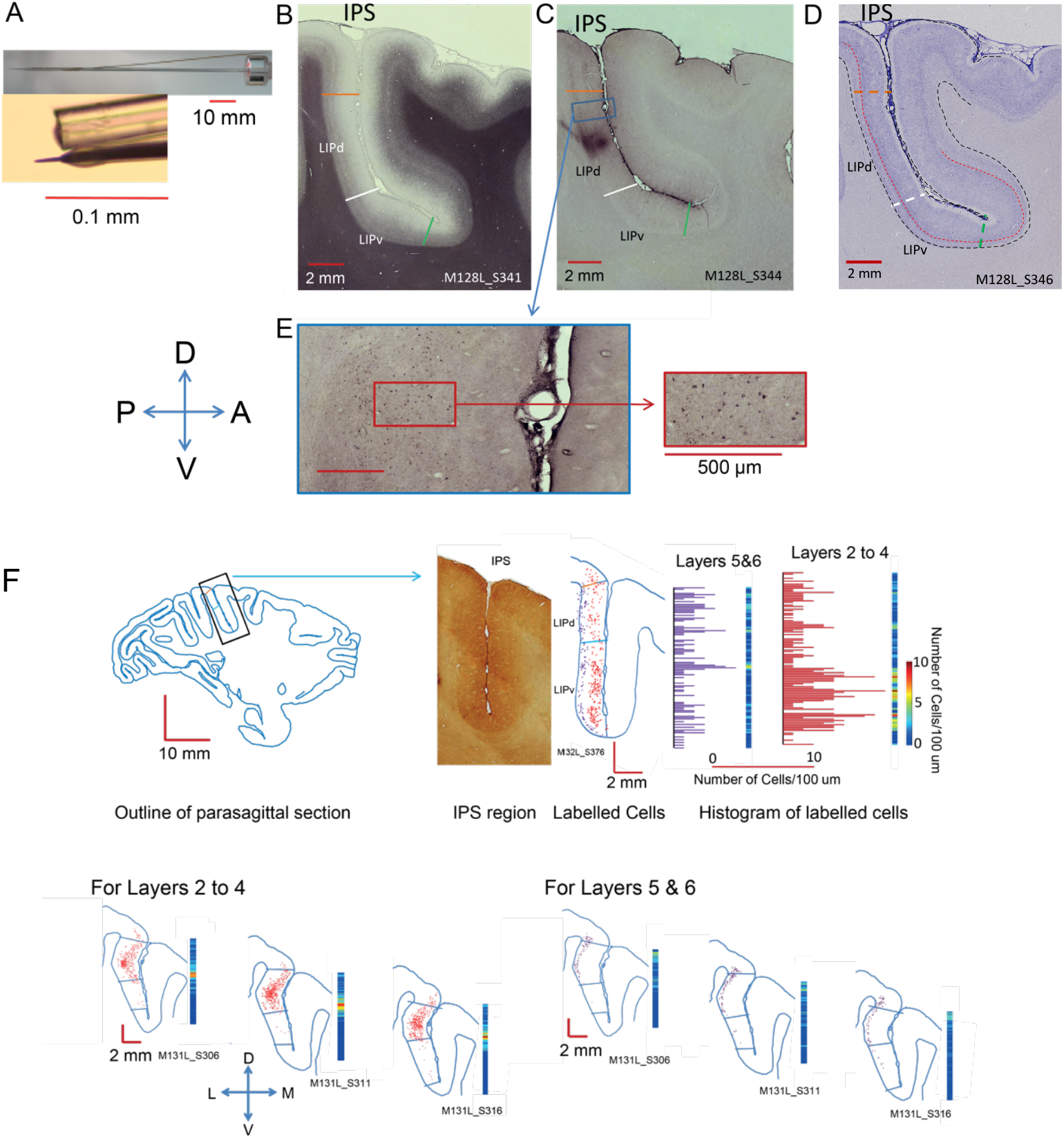
Tracer injections and analysis. A. Injections were placed using a tungsten electrode for recording neural activity, which was glued to a glass capillary tube filled with tracer [28]. **(B-D)** Parasagittal sections through the intraparietal sulcus. **(B)** This section was stained for myelin using the Gallyas method. LIPv is identified histologically by the more dense, extended intracortical myelin. **(C)** Nearest parasagittal section with CTb injection site in LIPd, taking a posterior approach; for LIPv, we took either a posterior approach (two animals) or a dorsal approach (one animal). **(D)** Nearest Nissl-stained section identifying the border between layers IV and V. **(E)** Section stained for CTb shows the retrogradely labelled neurons. **(F)** Method for the transformation of data of CTb labelled cells from individual parasagittal sections into 3D maps of intrinsic LIP connectivity. Labelled cells were converted to density measures along the dorso-ventral axis, separately for supragranular (Layers 2-4) and infragranular layers (5+6). Then, data were aligned along the medio-lateral extent of LIP.

First, we analysed three brains (M128, M129, M131) with an injection placed in LIPd (Fig. 2). A single injection point in area LIPd labels neuronal cell bodies throughout the medio-lateral and dorso-ventral extent of LIPd itself (Fig. 2A). The extent of labelled neurons in LIPd is comparable to the long range of intrinsic connectivity within extrastriate visual areas V5/MT or TE [28, 29]. In contrast, only a single cluster of neurons in LIPv projects to the injection site in LIPd. We analysed the three-dimensional pattern of labelling by projecting the labelled cell data from each section onto a line, separately for upper and lower layers (Fig. 1F). The labelled cell densities were then aligned across the parasagittal sections using landmarks and area borders. Labelling tends to be stronger in the dorso-lateral plane of the injection site and medial to the injection site (Fig. 2B). But the density of labelled cells drastically decreased near the LIPd/LIPv border as identified on neighbouring sections. The observation of a single cluster in LIPv projecting to LIPd is confirmed by the data from all three brains, regardless of the retrograde tracer (CTb or FG) used. Like the bulk of the LIPd label, this cluster appears near the same dorso-lateral line (M128, M131) or immediately medial to the injection site (M129) and is more pronounced in the upper layers. This pattern demonstrates a topographic, point-to-point input from LIPv to LIPd. This is the type of sparse, ordered projection one would see in early to mid-level sensory areas in primates, as for example from V1 to V2 or to V5/MT, and which underpins the neurophysiological responses in these areas [30-33].

**Fig. 2.**
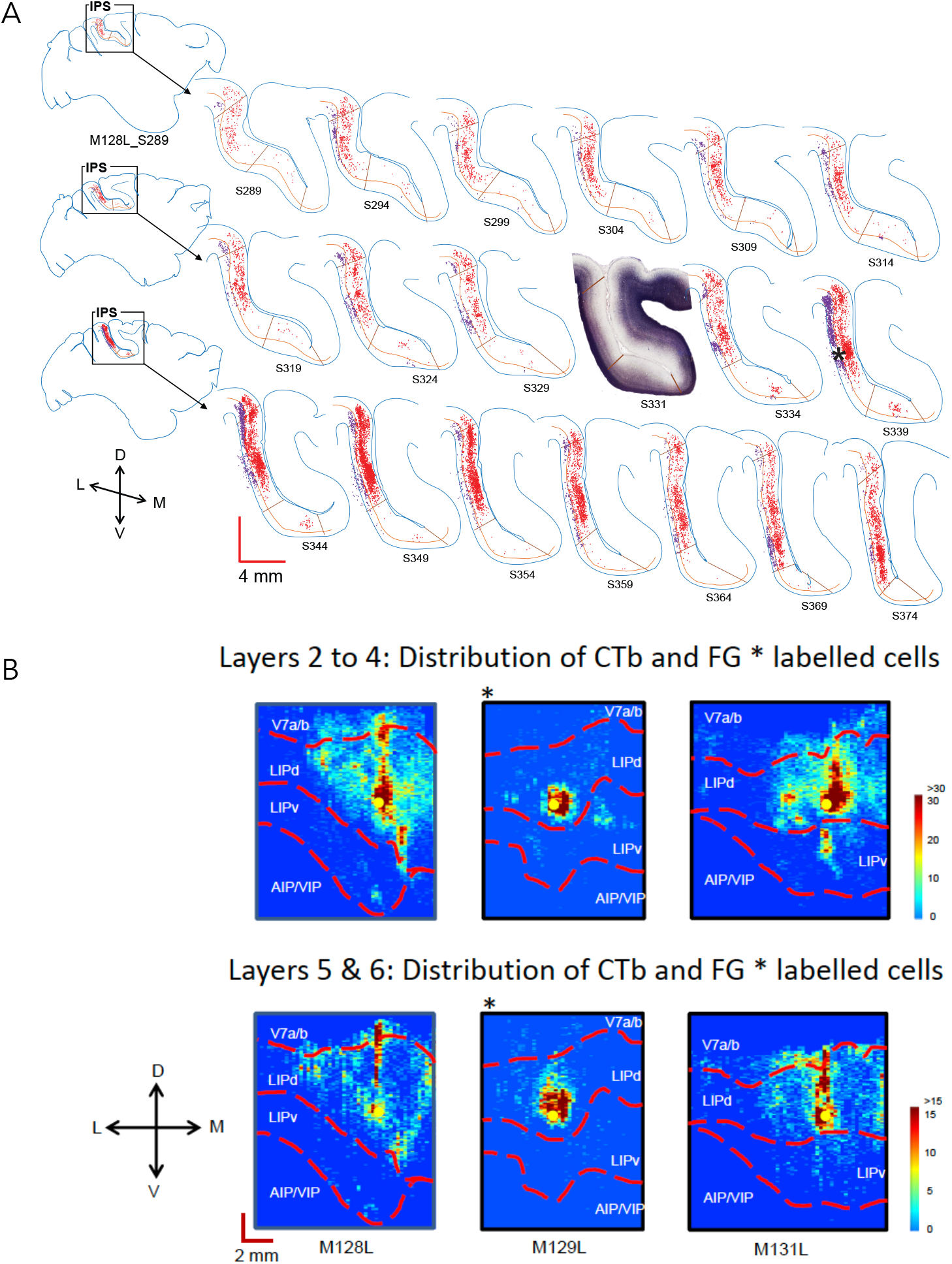
Neurons in LIPd receive topographic input from a single site in LIPv but widespread input from within LIPd. **(A)** A one-in-five series of parasagittal sections shows the distribution of labelled cells after a CTb injection into LIPd in the left hemisphere of one animal (M128L). Retrogradely labelled neurons can be found throughout the layers and the extent of LIPd, but there is only one clear cluster of labelled cells in LIPv that project to the injection site. The densest label in LIPd is medial to the injection site. The inset section shows the myelin definitions of LIPv and LIPd from an alternate series. Red cells are in layers I-IV, purple cells in layers V-VI; *– injection site. **(B)** The figures summarize the 3D-pattern of label (dorso-ventral; medio-lateral; supragranular and infragranular layers) across all three animals with a tracer injection into LIPd (denoted as yellow dot). M129 (*) received an injection of the retrograde tracer Fluorogold; M128 and M131 of CTb. All three animals show an LIPd-intrinsic, wide-spread network of neurons that project to the injection site in area LIPd itself. They also all show a single cluster of labeled cells in LIPv indicative of a topographic input from LIPv to LIPd. Legend cells/100μm. L – lateral, M – medial, D – dorsal, V – ventral.

### LIPv neurons receive projections from multiple distinct cell clusters in LIPd

In three further animals, we either placed single CTb injections into LIPv (M132, M140) or at the border between LIPd and LIPv (M127). The pattern of label shows a single point in LIPv received inputs from neurons all across LIPv, with the highest density of labelled neurons found around the injection site (Fig. 3A). There was also evidence of strong connectivity along the dorso-ventral axis in line with the injection site. In contrast to the topographic input from LIPv to LIPd, LIPv received more widespread inputs from several clusters of labelled cells across the medio-lateral extent of LIPd (Fig. 3B), mapping many points across LIPd to one in LIPv. The injection site at the border between LIPv and LIPd showed a similar result (M127). Thus, the LIPv-LIPd connectivity forms a circuit that potentially can map one location in topographically organized LIPv to one of many others in a recurrent network. The functional nature of the LIP map(s) is under debate [27]. Next, we investigated this at the level of single LIP neurons.

**Fig. 3.**
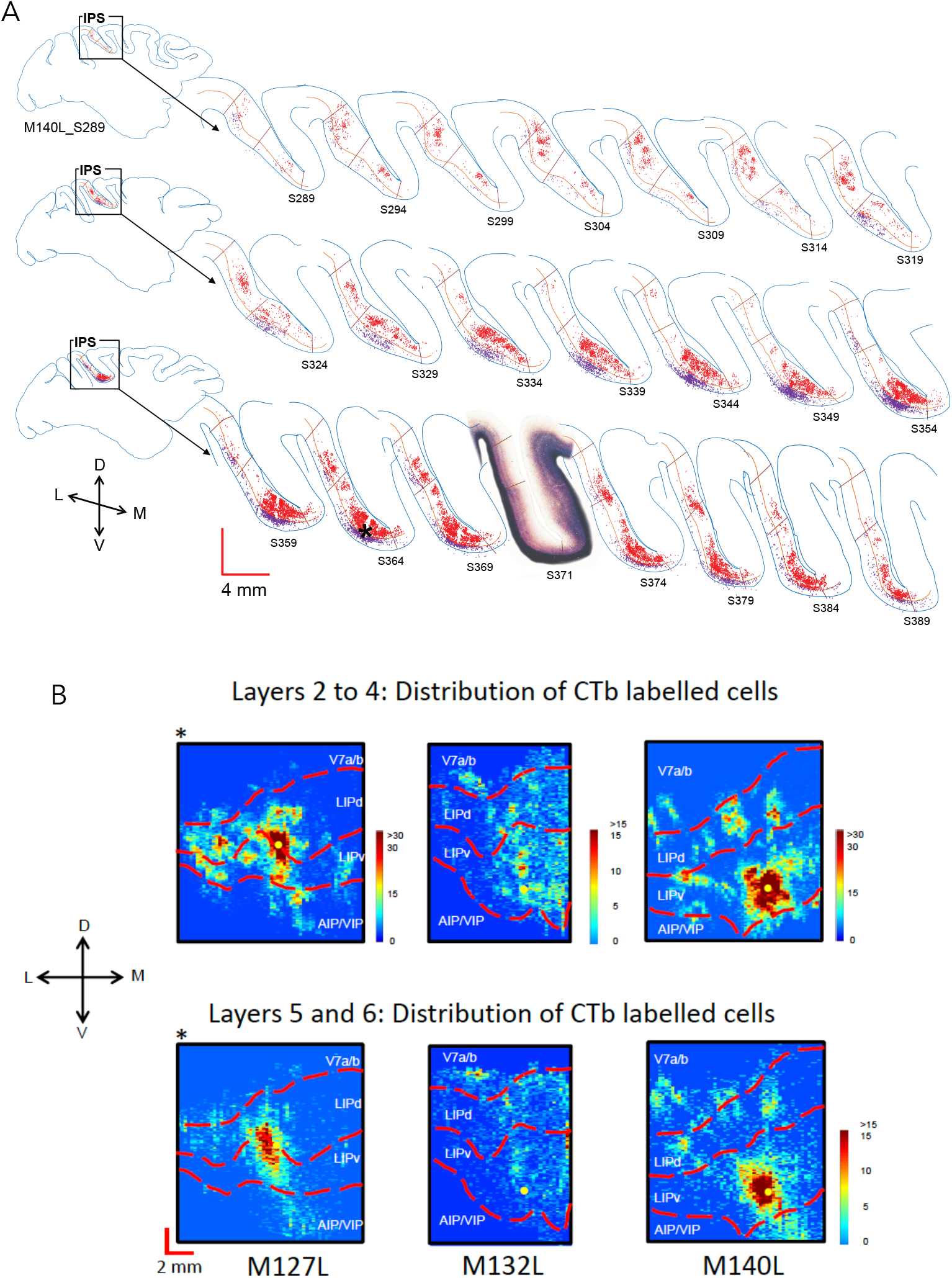
Neurons in LIPv receive wide-spread input within LIPv and from several cell clusters in LIPd. **(A)** A one-in-five series of parasagittal sections shows the distribution of labelled cells following a CTb injection in LIPv (M140L). Retrogradely labelled neurons can be found in supra- and infragranular layers across LIPv. In LIPd, several clusters of labelled cells are seen throughout the medio-lateral extent of LIPd. The inset section shows the myelin definitions of LIPv and LIPd from an alternate series. Red cells are in layers I-IV, purple cells in layers V-VI; *– injection site. **(B)** Figures summarize the 3D-pattern of label across all three animals with a tracer injection into LIPv (at the LIPd/v border for M127L). All three animals show an LIPv-intrinsic, wide-spread network of neurons that project to the injection site and multiple clusters of labeled cells in LIPd projecting to LIPv. Legend cells/100μm. L – lateral, M – medial, D – dorsal, V – ventral.

### LIP neurons have spatially separate motor response and sensory receptive fields

LIP combines both sensory and motor inputs from different sources with distinct functional topographic maps. We investigated how these maps might intersect in single neurons. We tested the neurophysiological response maps for 111 LIP single neurons in two macaque monkeys (41 from M133, 70 from M134), for which we could determine online a motor response field (MF) and a visual receptive field (RF). We systematically mapped their MF with a delayed saccade task and their visual RF with moving random dot (RDS) stimuli (Fig. 4). After carefully isolating single units offline and statistical analysis of stimulus und task related responses, 66/111 LIP neurons had a clear sensory receptive field (RF) (ANOVA/t-test p< 0.05), 60/111 LIP neurons showed a robust preference for a target location during the delayed saccade (MF) (ANOVA p< 0.05) and 45/111 LIP neurons had a significant RF and MF. Many neurons also showed clear tuning for direction of motion or binocular depth in their RFs (*data not shown)*. The distribution of RF and MF spatial response properties on their own matched those previously reported, with a preference for contralateral RFs but ipsilateral and contralateral MFs more evenly distributed [9, 18, 26, 34]. In many cases of cells with both RF and MF, we found the two were not congruent and could be even in opposite hemispheres, with RF and MF centres as far apart as 18 degrees (Fig. 4A-C). Thus, single LIP neurons could associate responses to a visual stimulus in one part of the visual field and planning a saccade to a distant location.

**Fig. 4.**
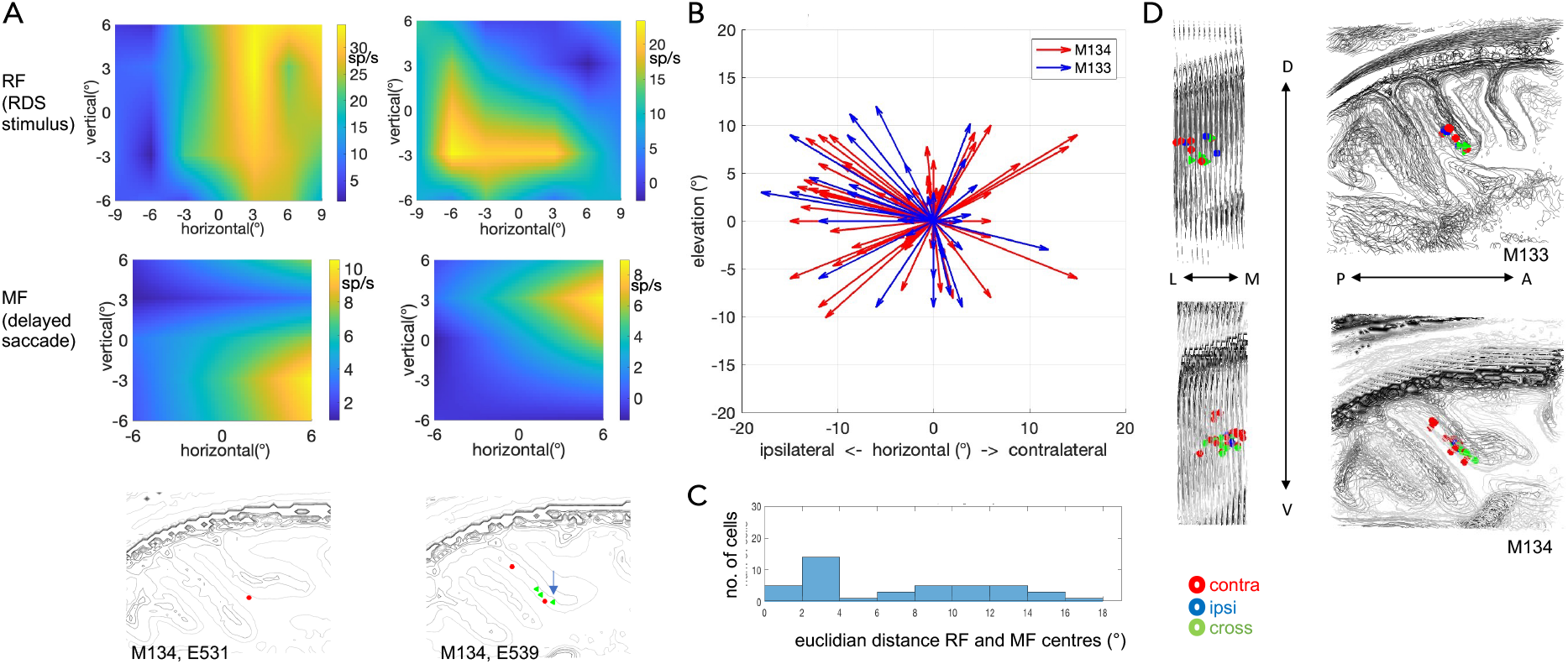
LIP neurons with spatially distinct RF and MF and their distribution. **(A)** Two example cells (columns) show spatially distinct visual receptive fields (RF) (top) and motor response field (MF) (middle) in a delayed saccade task. Scale is relative to central fixation at 0,0. Below is the location of the recorded neuron reconstructed on the animal’s structural MRI. **(B)** Direction and distance of MF centres from their corresponding RF centre (projected to 0,0), for all LIP cells, we could map qualitatively online (n=111). Some are very close to each other, others quite far removed. As most RFs are found in the contralateral visual hemifield and about half of the MFs on the ipsilateral side, many arrows point that way. Overall, a large range of different visual field associations can be found. **(C)** Distribution of absolute distances between RF and MF centres for LIP neurons with a significant RF and MF (n=45). In our sample, many LIP neurons have overlapping RFs and MFs but a considerable fraction can be as far apart as 12-18 degrees. (**D)** Spatial location of all LIP neurons with significant RF and MF for the two monkeys (M133, M134). LIP neurons with RF and MF across the two hemifields tend to be located more ventrally.

Stereotactically placed vertical penetrations into area LIP and careful records of the gray matter-white matter boundaries for each approach allowed us to reconstruct recording site of LIP cells in structural MRIs from the individual animals (Fig. 4D). These reconstructions indicate a bias for neurons with RF and MF in opposite hemisphere to be found more ventrally in our recordings.

## Discussion

We show a distinct, asymmetric pattern of connectivity forming a loop between anatomically defined ventral LIP (LIPv) and dorsal LIP (LIPd). LIPv provides a sparse, point-to-point projection to LIPd, like one would see in early or mid-level sensory areas. In turn, multiple clusters of LIPd cells project to one point in LIPv, linking potentially many points to one map location. Neurons within either sub-compartment of LIP are also widely interconnected. This circuitry links the distinct sensory, motor and cognitive inputs coming into LIPv and LIPd. Neurophysiological recordings demonstrated that the association of distinct sensory and motor planning signals is computed in single LIP neurons, which can have spatially distinct visual RFs and saccadic MFs. In sum, the neuroanatomical findings underpin a functional sensorimotor architecture that can link topographically organized, instructive visual stimuli to oculo-motor, attention or decision targets.

The most striking feature of the intrinsic connectivity is the sparse, topographic input from LIPv to LIPd in stark contrast to the reciprocal, wide-spread connectivity from LIPd and the widespread, within-compartment connectivity. This anatomical wiring patterns clearly distinguishes two operational compartments. The sparseness of the LIPv to LIPd projection is similar to that of early and mid-level visual areas and would be suitable to maintain a spatial map of visual field positions for visual stimuli and saccade targets. This finding is consistent with prior evidence of a topographically organized LIPv [9, 25, 26] and an oculo-centric representation of a perceptual decision formation [35]. Putting these anatomical findings into context of the cortico-cortical connectivity pattern of LIP [9, 11-15], we propose an operational circuit that can link spatial and object visual information received from V5/MT and V4 to saccade and attentional targets through the circuitry with FEF (Fig. 5). We have here potentially two recurrent network circuits: one intrinsic to LIP between LIPv and LIPd and another one looping through FEF. FEF could convey attentional or oculomotor planning signals via a feedforward-type connection into LIPd and also receives input from topographically organized LIPv. Further experiments are required to investigate dynamic changes in these circuits during shifting attention, saccade planning, perceptual decision-making and visual categorization tasks.

**Fig. 5.**
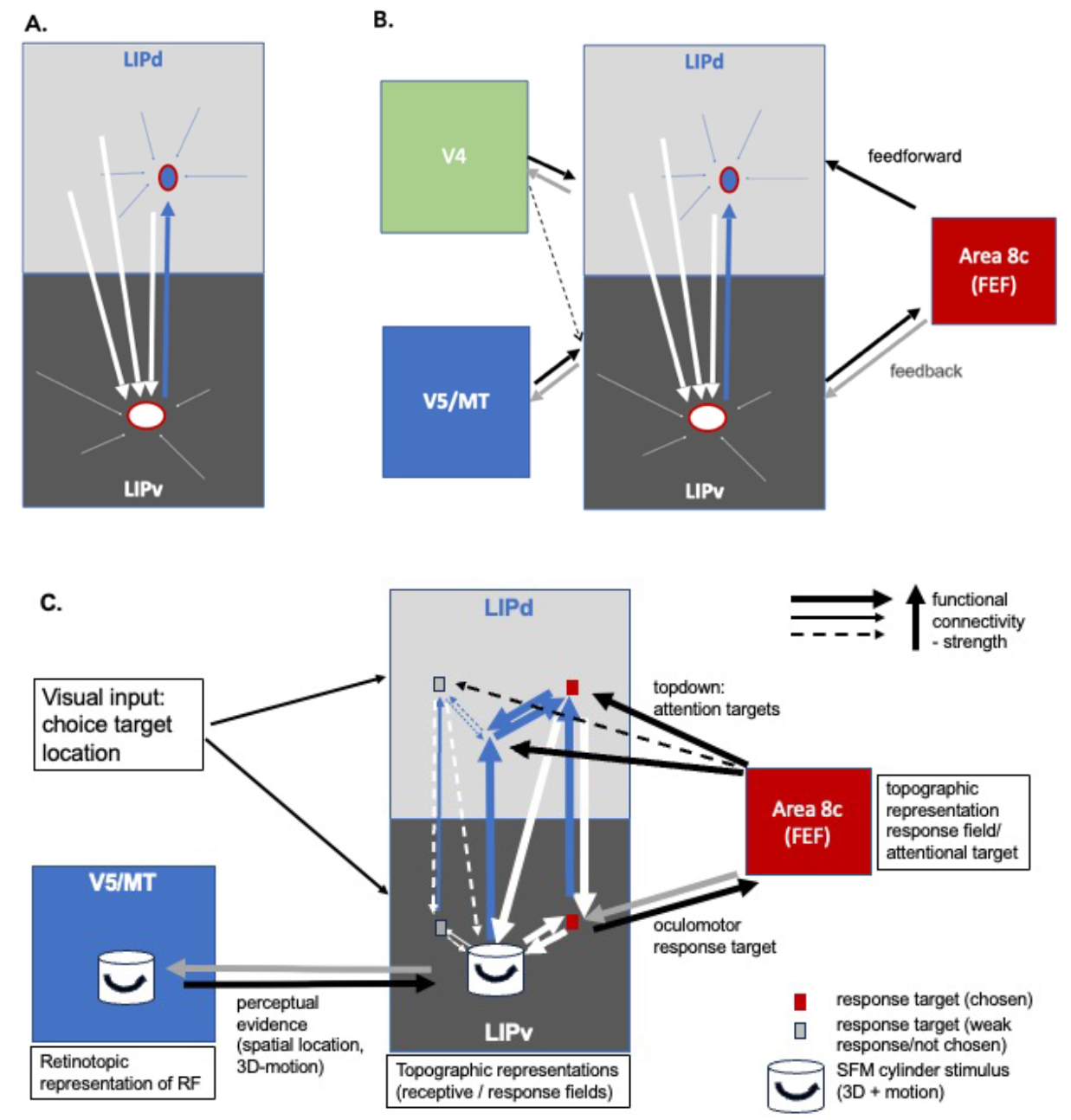
Summary of LIP intrinsic connectivity, (selective) wider network and proposed circuit for processing perceptual decisions about 3D motion in macaques. **(A)** This diagram summarizes the key findings of the intrinsic connectivity pattern between LIPd and LIPv from this study. LIPv, previously shown to be topographically organised [25], sends a point-to-point projection to LIPd. In turn, each point in LIPv receives inputs from a number of different regions within LIPd. **(B)** Illustration of selected, previously established cortico-cortical in- and outputs to LIPv and LIPd. Extrastriate visual area V5/MT projects exclusively to LIPv, not LIPd, while ventral stream visual area V4 has a stronger projection to LIPd [9, 13]. The frontal eye fields (FEF) send a feedforward-type projection to LIPd, and a feedback-type projection to LIPv [15]. **(C)** Illustration of the proposed processing scheme for perceptual decisions about 3D structure-from-motion stimuli, based on input of perceptual evidence represented in area V5/MT [37, 38] to LIPv. The intrinsic network between LIPv and LIPd can map one input-to-many locations. This might be achieved gradually in a recurrent network. Interconnected with a feedforward connection to LIPd and from LIPv, FEF signals convey an attentional shift from stimulus to choice target and thus facilitate the association of the stimulus with a specific choice target. The specific associations of RF and MF in single neurons we pick up are the product of the daily visual experience of the animals directing saccades to salient visual features and the specific training the animals underwent to learn to make perceptual decisions about 3D structure-from-motion objects.

The anatomical circuitry is reflected at the cellular level in the neurophysiological properties of the LIP neurons we mapped. Visual, attentional or saccade planning selectivity of single neurons in area LIP has long been described in a number of papers [9, 18, 23, 36], though usually individual studies put the emphasis on different properties measured in different behavioural contexts (delayed saccade for MF and fixation or anesthetized for RFs). The most common link between sensory RF and intentional MF is the reported response to a visual stimulus placed within MF, describing RF and MF as overlapping (e.g. [7]. Using stereotactically targeted, single neuron recordings, we show now that this is one of a range of spatial configurations and single LIP neurons can have a spatially distinct sensory RF from the MF.

The demonstration of spatially separated RFs and MFs for the same neurons offers a simple mechanistic explanation for seemingly contradictory studies in their interpretation of the role of area LIP in sensorimotor processing and in decision-making [7, 18, 36, 39, 40]. The neuronal responses and task contribution of LIP neurons would differ depending on how first, an LIP neuron was characterised and, subsequently, an instructive visual stimulus and the saccade targets are placed in relation to the RF and MF. Some of the consequences have been recently illustrated in an inactivation study by Zhou and Freedman [7] for overlapping RFs and MFs. We show now that there are other sets of neurons with different spatial configurations. Using stereotactical penetrations, we sampled LIP neurons for their responses to moving visual stimuli *and* delayed saccade responses, which we encountered regularly when recording in LIP. For those neurons we could map, about half had spatially separated RFs and MFs.

Activity of LIP neurons has been linked to cognitive functions, like perceptual decision-making about random motion dot stimuli [40, 41] and categorisation of such visual stimuli [7, 42]. Recent studies also suggest that the shifting perceptual evidence or attentional signals in such task are dynamically represented within LIP [6, 35, 43]. These data can potentially be explained by a common circuitry framework that dynamically associates relevant sensory cues with disparate locations representing upcoming attentional or saccade targets. Our anatomical and neurophysiological results delineate such a neural circuit that can flexibly link visual object information to a cognitive or behavioural target.

## Methods

### Animals

Combined neurophysiology and behavioural experiments were performed in two adult male macaque monkeys (*macaca mulatta*, M133, M134) weighing 13-17 kg. For each animal, structural MRI scan (MPRAGE; blackbone) were obtained under general anaesthesia. Based on the MR scans, animals were implanted with a titanium fixation device (Graymatter, USA) and a stereotactically placed, vertical titanium recording chamber (Thomas RECORDING GmbH, Germany) was mounted over the craniotomy above posterior parietal cortex to allow recordings parallel to the parasagittal/coronal planes.

For the histological study, six adult macaque monkeys (*macaca mulatta;* 4 females, 2 males, age range 4.8 – 12.4 years; weight 6.2 – 14.1 kg) were included (Table 1). All procedures were carried out under general anaesthesia. For each animal, a structural MRI scan (MPRAGE) was obtained. Five animals received a craniotomy and an MR-compatible chamber (Rogue Research Inc., Canada) centred on the lunate sulcus of the left hemisphere in a separate procedure prior to the injection. All procedures conformed to United Kingdom Home Office regulations on animal experimentation and to the European Communities Council Directives in force at the time.

**Table 1.** Details of animals and injections in the histological study.

| NHP ID | Sex | weight (kg) | Age (years) | Transport (h) | Tracer | Amount | Approach | LIP |
| --- | --- | --- | --- | --- | --- | --- | --- | --- |
| M127 | M | 14.1 | 12.4 | 113 | CTb (1%) | 120 nL | posterior | LIPd/v border |
| M128 | F | 6.2 | 4.8 | 186 | CTb (1%) | 80 nL | posterior | LIPd |
| M129 | F | 6.2 | 4.8 | 164 | FG (2%) | 80-100 nL | posterior | LIPd |
| M131 | F | 7.4 | 6.1 | 234 | CTb (1%) | 70-80 nL | posterior | LIPd |
| M132 | F | 6.3 | 6.3 | 278 | CTb (1%) | 80-100 nL | posterior | LIPv |
| M140 | M | 8.8 | 6.2 | 87 | CTb (1%) | 80 nL | dorsal | LIPv |

### Awake neurophysiology experiments and analysis

Monkeys M133 and M134 were trained to maintain fixation on a binocularly presented visual stimulus and to perform a delayed saccade task while their eye movements were video tracked (SensoMotoric Instruments [SMI], Germany). They received fluid rewards for correct responses or maintaining fixation and their daily fluid intake was controlled.

Visual stimuli were displayed binocularly using a Wheatstone stereoscope comprising of a set of mirrors and a pair of CRT monitors (Eizo FlexScan F78, UK) at a viewing distance of 84 cm. Mean luminance was. Monitors had a mean luminance of 42 cd/m^2^, a resolution of 1600 × 1200 pixels, a screen frame rate of 85Hz, and covered 26.7° × 20.1° of the subject’s visual field. Quadro K2200 (NVIDIA) video cards in multi-core Intel processor computers running Ubuntu 14.04 (Linux kernel 4.4.0-124-lowlatency) were used to drive the monitors. Stimuli were programmed using PsychToolbox (3.0.14) running in Matlab R2015b (The Mathworks, USA). All stimuli were presented on a mid-grey background, using OpenGL alpha-blending to obtain sub-pixel resolution.

Animals performed a fixation task or a delayed saccade task for mapping the motor response fields [34, 44]. For the fixation task, monkeys received a fluid reward for maintaining fixation on a central white fixation point (0.2° diameter circular dot) for 1s (RF mapping and direction tuning). If the gaze fell outside of the fixation window (radius 1°-2°), the trial was aborted. For all neurons, we mapped visual receptive fields with a patch of coherently moving black-and-white dots (0.2° diameter) while the monkey maintained fixation. The position, speed, direction, and circular size of the patch of moving dots on the computer screen was adjusted interactively to maximise neuronal responses of the isolated single unit. For a large fraction of LIP neurons (67 out of 111), we also mapped the receptive field quantitatively by probing the visual field with a circular patch of moving, black and white dots (dot diameter 0.2°; patch radius 3°; random motion or if little response also coherent motion). The patch was randomly presented at one of 4 y-axis positions (−6°, −3°, 3°, 6°) and 6 x-axis positions (−9°, −6°, −3°, 3°, 6°, 9°). For all neurons, we also measured direction tuning in the mapped receptive field using a patch of black and white, coherently moving dots matched in patch size and dot speed to the receptive field (randomly interleaved: directions of 0°, 60°, 120°, 180°, 240°, 300°, and blank screen), which we used to test for a significant response to the visual stimulus if no quantitative RF map was available (n = 44).

For the delayed saccade task: after fixating a white central fixation point (0.2° diameter) for 0.5s, a white saccade target dot (0.2° diameter) appeared in the periphery. The saccade target was randomly presented at one of 5 y-axis positions (−6°, −3°, 0°, 3°, 6°) and in either 2 or 7 x-axis positions (−6°, 6° or −9°, −6°, −3°, 0°, 3°, 6°, 9°) from the centre of the screen. When the fixation point was extinguished after 1s (“go” signal), the animal was required to make a saccade to the target (radius 1.5°, 2.0° or 2.5° [2 sessions]) within 300 ms to obtain a fluid reward.

Neurophysiological recordings in area LIP were carried out through a vertical chamber using a smokestack with guide grid (1mm spacing; Crist Instruments Co Inc, USA), which was centred stereotactically over LIP using MRI guidance (Brainsight, Rogue Research Inc.). This allowed us to systematically map neural response properties along dorso-ventral penetrations in a circular window with a diameter of 14 mm). We recorded single units with tungsten microelectrodes coated with polyimide tubing (0.8 –1.2 MΩ impedance at 1 kHz; FHC, MicroProbe Inc., USA) inserted into the cortex through a short guide tube and advanced using a hydraulic microdrive (MO-97, Narishige Group, Japan). Electrical signal were filtered, amplified, and displayed through visual and audio monitors, and stored to computer disk (Blackrock recording system, Blackrock Microsystems LLC, USA). Binocular eye movements of the animal were recorded with an SMI eye tracker and the iView X system program (SensoMotoric Instruments [SMI], Germany). Offline spike sorting was carried out with Blackrock Offline Spike Sorted (BOSS) (Blackrock Microsystems LLC, USA) and were subsequently analysed using software tools developed in MATLAB (MathWorks, USA).

Area LIP was identified by carefully recording electrode depth and gray and white matter boundaries every 200 μm on each penetration. These were aligned by x- and y-position on the guide grid to the structural MRI scans taken from the animals before chamber implantation. First, we checked the fit of penetrations for each parasagittal plane, subsequently also in the horizontal plane across all penetrations. We found the best fit for the neurophysiological data to the MRI structural scan for each monkey by adjusting the overall positioning of the centre of the guide grid over cortex by 3-5 mm in the horizontal plane. This was necessary due to potential alignment changes during the surgery to fit the custom-made Titanium chamber (Thomas Recording, Germany) to the skull. Once aligned, the absolute depth of individual penetrations could be adjusted by 1-3 mm (necessary due to the use of different lengths guide tubes for penetrating the dura). Recorded gray matter – white matter boundaries of each penetration and horizontal relationship between penetrations was preserved as recorded. Response properties on all included penetrations showed at least qualitatively a clear delayed saccade response in the target structure.

To establish the sensory receptive field, we tested for significant variation in evoked response to the presented stimuli in the quantitative mapping task (after deducting firing rate during fixation without a stimulus; ANOVA, p <0.05). If there was no quantitative RF mapping collected, we used the responses from the direction tuning measurements, comparing RDK stimulus trials against blank screen (t-test p<0.05). To examine firing rates for the delay period, we subtracted from the firing rate for the delay period the firing rate of the preceding fixation period and tested whether there were significantly different responses for the different saccade target locations (ANOVA, p<0.05). We identified the saccade target region that led to the strongest responses during the delay period as the saccade response field of the cell. To smoothly map the mean firing rate across the sampled visual space, we used Matlab commands (*meshgrid, interp2, surf*) to linearly interpolate between the center positions of sample stimuli. The resultant 3D surface is viewed from above, with the colour scale indicating mean firing rate above baseline as describe dabove.

### Tracer injections

Tracer injections were placed either in a recovery procedure or in a five-day terminal procedure, all under general anaesthesia. For induction, animals received ketamine (7.5mg/kg), midazolam ([Hypnovel], 0.1mg/kg), xylazine (0.125mg/kg (i.m.) as well as atropine sulphate (0.05mg/kg i.m.). Animals were intubated and artificially ventilated (2-3% Sevoflurane). The head was placed in a stereotaxic frame. Non-invasive blood pressure, heart rate, electrocardiogram, oximetry, and body temperature were monitored throughout. Hartman’s solution (Ringer’s lactate solution) was given through a cannulated saphenous vein. For the animal without a chamber, a craniotomy was made. For the visual stimulation, contact lenses of +3D were placed on both corneas and spherical lenses placed to focus the eyes on a screen at 1.14 m.

Under general anaesthesia (∼2.5% Sevoflurane by inhalation or 0.006 mg/kg/hr Sufentanil i.v. + 0.5% Isoflurane by inhalation), we made a small durotomy and advanced a glass capillary tube containing the retrograde tracer attached to a tungsten electrode (Microprobe Inc., impedance 0.65 - 0.75 MOhms) (Fig. 1A). Trajectory and target site for the injection were identified by a combination of MR-based guidance (Brainsight, Rogue Research Inc., Canada), stereotactic coordinates and neuronal recordings on approach. For LIPd injections, a posterior approach was chosen to insert the electrode into cortex with an angle of 5 – 20° over the horizontal plane 1-2 mm anterior to the lunate sulcus and 10 – 11mm lateral to the midline. For LIPv we chose either a posterior or a dorsal approach entering anteriorly to the intraparietal sulcus (IPS). We recorded graymatter/whitematter boundaries as we advanced the electrode and mapped multi-unit visual receptive fields.

Five monkeys (M127, M128, M131, M132, M140) were injected with 70-120nl retrograde tracer Cholera Toxin subunit β (CTb, 1% low salt solution, List Biological Labs, USA), one female monkey (M129) was injected with 80-100nl retrograde tracer Fluorogold (FG, Fluorochrome LCC). After 87-278 hours, animals were perfused and the tissue processed.

### Histology

After transcardial perfusion with 0.9% heparinised PBS and 4% Paraformaldehyde the left hemispheres were removed and cryoprotected in 10%, 20% and finally 30% sucrose in 0.1M Phosphate Buffer. Tissue was sectioned parasagittally at 50 μm on a freezing stage microtome. A one-in-five series each were stained for Gallyas [45] and Nissl [46] to determine the boundary between LIPd and LIPv and to adjacent areas. Nissl-stained sections were also used for discrimination between different cortical layers. Another one-in-five series was reacted for CTb staining or FG staining. All reactions were carried out on free-floating sections to maximise penetration. If reactions were not carried out immediately, they were stored in cryoprotectant at −20°C. Sections were treated with a peroxidase blocker to block non-specific sites. Next, sections were incubated for 16-20h at 4°C in peroxidase blocker containing either primary antibody against cholera toxin b subunit (List Biological laboratories) diluted 1:9000 for sections stained for CTb or rabbit anti-fluorogold (Fluorochrome LCC) diluted 1:1000 for sections stained for FG. This was followed by incubation for 90min at room temperature (RT) with secondary antibody anti-goat or anti-rabbit IgG Biotin (Sigma) diluted 1:400 or 1:300, respectively. All sections were treated with ABS (Vector Elite) for 60min at RT before they were reacted with SigmaFast™ DAB for 5min at RT to visualise stained cells. Finally, sections were mounted on gelatinised slides, air-dried at 37°C, dehydrated and cover-slipped with DPX (Merck).

### Histology analysis

We quantified the distribution of retrogradely labelled cell bodies in LIP using a computerized microscope (Neurolucida, Microbrightfield Ltd; Matlab, MathWorks) in histological sections. The outline of the intraparietal sulcus and the grey matter/white matter boundary were drawn at 2x magnification. Locations of CTb or FG labelled cells were registered by marking cell bodies at high magnification of x10. FG- and CTb-labelled cells showed distinct patterns of label in the cell body allowing good identification. We analysed a complete one-in-five series of parasagittal sections through LIP for each monkey. The most lateral and medial sections which only showed a low number of labelled cells or in which border discrimination of LIP was not possible were excluded from the quantitative analysis. For one hemisphere stained for CTb (M127; injection at the LIPv/d boundary), the injection site included layer I leading to a high number of labelled cells in layer I. These cells were excluded from analysis in order to compare the labelling patterns across hemispheres.

All injection sites were neurophysiologically identified at the time of injection. The effective tracer up-take zone is thought to be the darkest area around the injection site in which labelled cells are not discernable under bright illumination and high magnification (x10)[47]. Such a core injection sites could be confirmed in all but one animal histologically (M132), where the visible track would have entered between the sections of the stained series. Injection sites were usually small, around 500-800 μm in diameter and well circumscribed with some lighter diffusion of tracer around it. We have previously confirmed that CTb is not taken up by fibres of passage [28], so the presented results show specific gray matter connectivity.

Interleaved sections stained for Gallyas or Nissl were examined for changes in myelination and cell patterns in LIP and adjacent areas at a magnification of 10x to discern borders between areas and subdivisions of LIP. In most cases the investigation of nearby Gallyas sections were sufficient to discern the border between LIPd and dorsal area 7a. LIPd showed two prominent bands of Baillarger and lighter myelination than 7a [9-11]. In the few sections, in which this criterion was not clear, we also examined Nissel-stained sections for differences in cell densities and cell arrangements between 7a and LIPd [9, 11]. The border between ventral and dorsal LIP was readily distinguishable by differences in myelination patterns [9-11] (Figure 1B). The discrimination of the border between LIPv and the adjacent ventral intraparietal area (VIP) was based on the decrease in myelination in VIP. In some sections, this was supplemented by examining Nissl-stained sections for differences in cell densities and organization [10].

The border between layer IV and layer V was outlined based on differences in cell density and appearance between layers in Nissl sections interleaved with the CTb/FG labelled sections using Neurolucida at 10x magnification. These outlines were overlaid on the CTb and FG labelled sections to determine the border of layers IV/V. In sections stained for CTb layer IV hardly contained labelled cells which made it easier to define the boundary to layer V.

To examine the pattern of labelled cells in across sections, we created cell density maps (Figure 1F). We then normalised the cross-section scale bar to the highest density of labelled cells found within LIP in this hemisphere. We analysed supragranular layers (ii-iv) and subgranular layers (5-6) separately and aligned the 2D plots across LIP sections.

## Acknowledgments

The authors thank Greg Daubney for the excellent histology, the Oxford BMS staff for expert care of the animals, and Andrew Parker for comments on the manuscript. This research was supported by German Research Foundation (DFG) grant KR 5138/1-1 & KR 5138/3-1 (Heisenberg Professorship) (KK), German Research Foundation (DFG) grant SFB1436 project C05 (KK), Biotechnology and Biological Sciences Research Council (BBSRC) grant BB/H016902/1 (KK), Wellcome Trust Strategic Award 101092/Z/13/Z (KK), and a Royal Society (UK) University Research Fellowship (KK).

